# Feedforward attentional selection in sensory cortex

**DOI:** 10.1101/2022.06.06.495037

**Authors:** Jacob A. Westerberg, Jeffrey D. Schall, Geoffrey F. Woodman, Alexander Maier

**Affiliations:** Department of Psychology, Vanderbilt Brain Institute, Vanderbilt Vision Research Center, Vanderbilt University, Nashville, TN 37240, USA; Centre for Vision Research, Vision: Science to Applications Program, Departments of Biology and Psychology, York University, Toronto, ON M3J 1P3, Canada

## Abstract

Salient objects stand out (pop-out) from their surroundings, grabbing our attention. Whether this phenomenon is a consequence of bottom-up sensory processing or predicated on top-down influence is debated. We show that the neural computation of attentional pop-out is embedded in the earliest cortical sensory response, seemingly void of feedback from higher-level areas. We measured synaptic and spiking activity across cortical columns in mid-level area V4 of monkeys searching for an attention-grabbing stimulus. Indexed by reaction times and behavioral accuracy, attention was captured at variable times. This moment of attentional capture occurred within the earliest feedforward response, both in terms of timing and spatial location. Moreover, the magnitude of the earliest sensory response predicted reaction times. Crucially, errant attentional selection and consequent behavior was associated with errant selection in sensory cortex. Together, these findings demonstrate a dominant role for feedforward activation of sensory cortex for dictating attentional priority and subsequent behavior.

**In brief:** Why do certain objects stand out from their surroundings and seemingly grab our attention? In this study, Westerberg et al. determine that attentional selection for salient objects in our environment is computed in sensory cortex as soon as sensory information arrives.

**Highlights:**

- Early sensory responses in V4 predict attentional selection and behavioral responses
- Errant attentional selection in sensory cortex precedes errant behavior
- Tonic modulation of sensory cortex can regulate attentional selection

## Introduction

We constantly filter through our complex environment to extract information pertinent to our goals. In this effort, some objects seem to grab our attention when they differ from their surroundings. However, the mechanisms supporting this attentional prioritization through salience-based capture (“pop-out”) remain elusive (Folk et al., 1992; Luck et al., 2021; Theeuwes, 1993). The behavioral and phenomenal consequences associated with pop-out are frequently described as “exogenous attention” or “stimulus-driven attention” (Chun et al., 2011), suggesting that feedforward sensory processes take a preeminent role. However, this has never been demonstrated directly. As a consequence, attentional prioritization through salience-based capture has been conflictingly theorized to arrive automatically and feedforward (Theeuwes, 1993) or via cognitive mediation (Folk et al., 1992). An intermediate hypothesis proposes that an automatic priority signal is generated in response to attention-grabbing objects which is biased to promote behaviorally useful objects (Luck et al., 2021).

We tested the predictions of these competing theoretical views by recording across the layers of primate area V4 while macaque monkeys searched for oddball targets in arrays of objects. Area V4 is ideal for testing predictions of the competing accounts of attentional capture as it receives from, and projects to, both earlier visual cortical areas and higher-order cortex implicated in cognitive control (Roe et al., 2012; Ungerleider et al., 2008) while showing robust attentional modulation (Luck et al., 1997) during visual search (Bichot et al., 2005; Ogawa and Komatsu, 2004).

We found that the neural computations underlying attentional capture occur within the earliest synaptic and spiking activation within the feedforward-recipient layers of V4 that comprise the initial spatio-temporal volley of sensory responses. This finding is incompatible with hypotheses involving extensive feedback from higher areas. Instead, these data suggest that “exogenous” or “stimulus-driven” attentional priority largely arises from bottom-up processes in early sensory cortex.

## Results

The feedforward account of attentional capture posits that attention-grabbing objects automatically engender capture (Luck et al., 2021; Theeuwes, 1993). For this account to hold, feedforward sensory activation elicited by these objects should predict behavioral measures of attentional capture, such as reaction time. This finding would suggest the priority (Fecteau and Munoz, 2006) of attention-grabbing objects is computed rapidly and in sensory cortex. Priority indexes the utility (as opposed to a sensory feature) of a stimulus. In pop-out, salience and priority are tightly coupled, yet these two attributes are experimentally distinguishable since a non-salient stimulus can sometimes be (erroneously) chosen as having the highest utility. However, feedforward sensory activation has not yet been shown to be tightly coupled to behavioral accuracy and reaction time in attention-demanding search tasks.

In contrast, the feedback account of attentional capture puts forward that stimulus features are prioritized more or less independently of the magnitude of sensory activation. In the extreme case, this view predicts modulation of cortical activity in the absence of visual stimulation. The latter phenomenon might manifest, for example, as persistent changes to ongoing activity, or during intertrial periods (Luck et al., 1997; Supèr et al., 2003; van der Togt et al., 2006). Top-down attentional modulations of neural activity can manifest when spatial selective attention is deployed (Armstrong and Moore, 2007; Bosman et al., 2012; Desimone and Duncan, 1995; Fries et al., 2008, 2001; Gregoriou et al., 2012; MartÍnez-Trujillo and Treue, 2002; Martinez-Trujillo and Treue, 2004; McAdams and Maunsell, 2000, 1999; Moore and Armstrong, 2003; Moran and Desimone, 1985; Nandy et al., 2017; Ni and Maunsell, 2019; Reynolds et al., 2000, 1999; Reynolds and Chelazzi, 2004; Reynolds and Desimone, 2003; Roelfsema et al., 1998; Treue and MartÍnez Trujillo, 1999; van Kerkoerle et al., 2017; Westerberg et al., 2021), but whether this feedback-driven mechanism of attentional modulation is also instantiated for pop-out selection is an open question. Finally, on the intermediate account we expect both mechanisms to emerge.

We first tested predictions of the feedforward hypothesis. If attentional capture were computed in a feedforward fashion, we expect two key pieces of evidence in V4 presented with an attention-capturing oddball (Figure 1A-B). For one, oddball signals should propagate through cortical pathways in a pattern specific to feedforward processing (Douglas and Martin, 1991; Douglas et al., 1989; Maier, 2013; Mitzdorf, 1985; Schroeder et al., 1998; van Kerkoerle et al., 2014). Second, the stimulus-driven response to the oddball (that is, the initial volley of activity preceding attentional feedback (Roelfsema et al., 2007)) should predict behavior. Both these tests require a neural measure of attentional capture. Here we do so by quantifying a time-resolved attentional capture (oddball selection) metric via population reliability analysis (Figure 1C-E). We also identified when the oddball most effectively captured attention by sorting trials by reaction time (Figure 1F).

**Figure 1.**
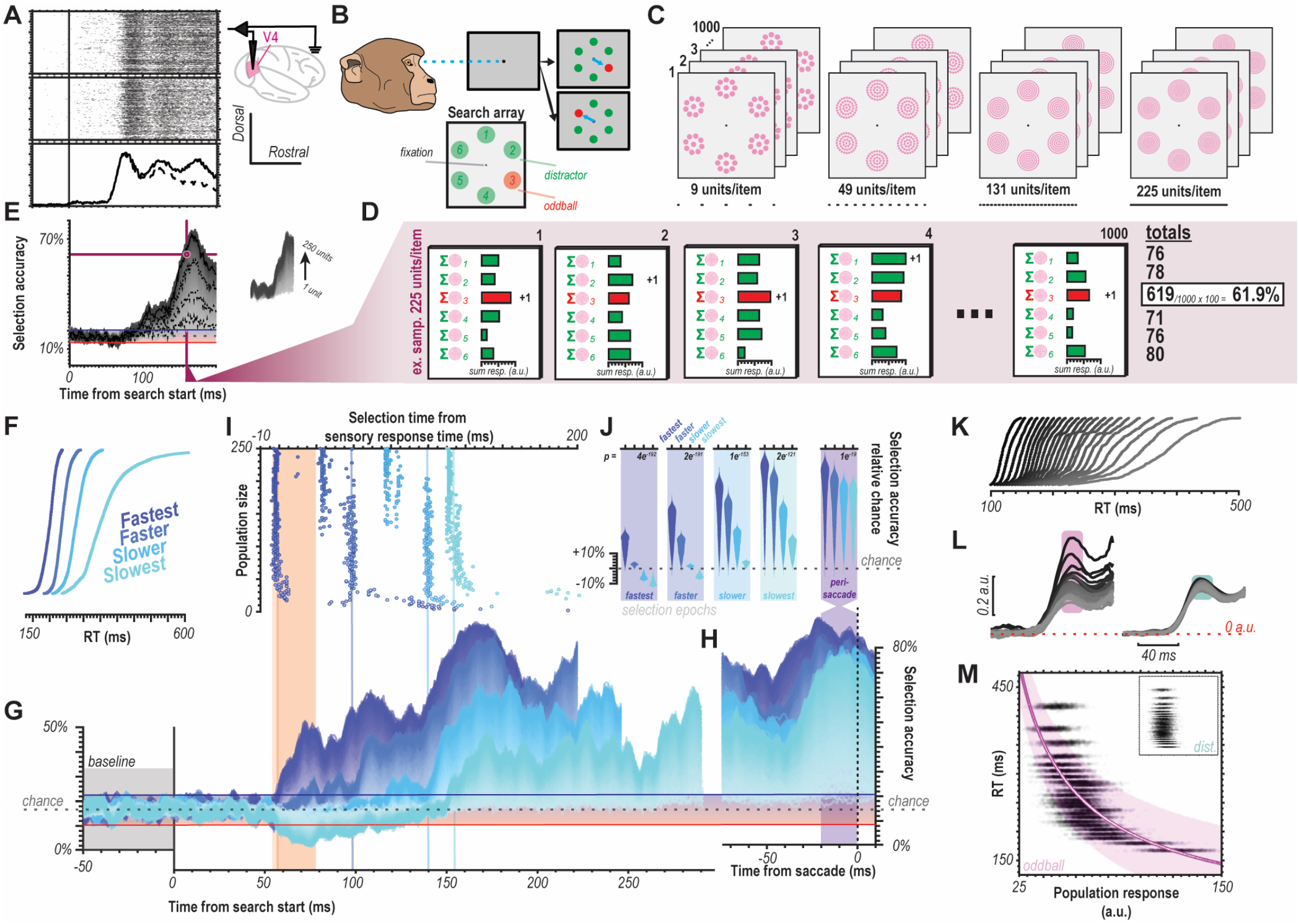
Temporal evidence for feedforward attentional selection in sensory cortex. **(A)** Neurophysiological recordings took place in mid-level visual cortical area V4. Top and bottom rasters show example of 1000 randomly chosen multiunit responses to an oddball or distractor stimulus, respectively. Traces at bottom show convolution of average multiunit responses to oddball (solid) vs. distractor (dashed) stimuli. **(B)** Monkeys performed 6-object color oddball search by making an eye movement to the oddball following presentation of the stimulus array. **(C)** Visualization of population reliability analysis, with magenta circles representing multiunits comprising the response to each stimulus. Four panels show example population sizes (9, 49, 131, 225) with stacking indicating the 1000 sampling simulations performed. **(D)** The summed multiunit response of each of the 6 (1 for each search item) samples was computed and their maximum was defined as selection of the associated stimulus. Each sample is a randomly selected set of trial-level multiunit responses for the corresponding stimulus type (i.e., oddball or distractor). This was repeated 1000 times for each population size for each millisecond across time to compute an oddball selection metric (percent of time oddball had the largest magnitude response across sampling simulations). **(E)** Oddball selection as a function of time in pop-out search for population sizes 1-250 considering all correctly performed trials (n=1320282 multiunit trial-wise responses) across both monkeys (n=2 monkeys, n=29 sessions). Values above the chance window indicate reliable population selection of the oddball. Chance is 16.67% for 6-object search. Chance window is the empirically measured variability in selection accuracy in the absence of visual stimulation as computed as the 99% confidence interval during the baseline period (100 ms prestimulus epoch). Four example traces highlighted as black dashed lines correspond to the 4 example population sizes show in C. Crossing point of magenta vertical and horizontal lines, denoted by the magenta circle, indicate time and population size combination exemplified in D. **(F)** Cumulative density functions (CDF) of reaction times (RT) organized in quartiles (n=2 monkeys, n=29 sessions). **(G)** Oddball detection for each RT quartile shown in F. Data clipped 10 ms prior to each respective median RT. Orange highlights duration of the initial transient of sensory response. Population sizes 1-250 for each bin represented as lightest to darkest traces. **(H)** Oddball selection across time for each of the 4 quartiles, aligned on gaze shift to the oddball. Population sizes 1-250 for each bin represented as the lightest to darkest traces. Chance window is identical to that in G. **(I)** Time when each population size for each bin first exceeded chance threshold in G. Color indicates data from each RT quartile. Orange highlight indicates the duration of the initial transient of sensory response. **(J)** 20 ms average oddball selection for each RT quartile immediately following the time where population 250 exceeded the chance window as well as the window immediately preceding behavioral reaction time (far right). Background highlight indicates each selection epochs’ corresponding quartile. Violin plots are relative to chance (16.67%). Statistic in upper righthand corner of each subpanel indicates result of ANOVA between quartiles. **(K)** CDFs for RTs divided into 24 bins. **(L)** Multiunit responses to oddball (left) vs. distractor (right) corresponding to 24-bin RTs. Darkest traces are fastest trials, lightest are slowest. Traces start at visual display. **(M)** Bayesian modeling 24-bin data with population size 250. RT as a function of population feedforward response (mean 58-78 ms post-display) to oddball, 1000 trials for each bin. Inset is distractor responses on identical scale. Magenta line shows result of power function fit. White data in magenta line are median estimates for simulated trials. Magenta cloud is 89% credible interval of median estimates for fit.

### Temporal evidence for feedforward selection

We first evaluated temporal evidence via population reliability analysis. Population reliability analysis has previously been employed to derive selection times in a decision-making task (Bichot et al., 2001). Based on neurophysiological data, this analysis provides quantitative insights into both when a selection process is completed as well as the neural population size that is required for this selection to occur reliably. Briefly, population reliability analysis assumes there is a neural population representing each of the selectable objects or surfaces in a task. At each millisecond in time, we summed the activity of a randomly chosen population of multiunits to determine which item produced the largest population response. As we did not simultaneously record all 6 stimulated regions of V4, we instead employed sampling simulations. That is, we subsampled responses across trials representative of the varying stimulation conditions (i.e., oddball vs. distractor inside receptive field). The stimulus yielding the largest response in this sample is taken as the selected item. This process is then repeated, each time taking randomly selected trial-level multiunit responses to the stimulus array to determine the frequency with which the oddball is selected. Crucially, this analysis can be performed over time to determine when population-level selection for the oddball stimulus is significant relative to chance. This analysis also affords the ability to vary the number of multiunits included in the population to determine the requisite population size to detect population-level selection. We defined the oddball selection frequency computed by this analysis as our attentional capture metric. Values exceeding chance threshold indicate reliable, statistically significant attentional selection of the oddball stimulus (see Methods and Figure legends for details on the statistical hypothesis tests).

To illustrate this analysis, consider an example calculation. First, we determined the frequency with which the oddball is accurately selected 130 ms after presentation of the array for a population size of 225 multiunits (Figure 1D-E). We assumed each of the 6 items is represented by the activity of 225 multiunits. We randomly selected 225 trial-level multiunit responses for each type of relevant stimulus. We summed those 225 multiunit responses for each item. The item with the largest summed response was tallied. We then repeated this process 1000 times using random (Monte Carlo) sampling. Of those 1000 samples, we counted the tallies for the oddball stimulus to find the frequency with which the oddball evoked the largest summed response (61.9% of the time, Figure 1D). That provided 1 data point on the ordinate in the selection time plots (e.g., Figure 1E). All other points were calculated by changing the time window (abscissa) and/or the number of multiunits (trace).

We found that oddball detection varied in concert with reaction time (Figure 1G-H). Moreover, significant detectable differences in population activation to oddball vs. distractor stimuli can already be observed in the earliest (50-60 ms following visual stimulation) sensory responses (Figure 1I). Crucially, we observed significant differences in the attentional capture metric across four successive epochs (bins) of the feedforward response (Figure 1J). This observation indicates that while the initial response to the oddball may not always evoke the largest population response, it entails sufficient information to predict the associated reaction time.

Next, we more finely investigated the exact relationship between population responses and reaction time (Figure 1K-L). No systematic differences in distractor responses were observed with reaction time; however, oddball response covaried nonlinearly with reaction time (Figure 1L-M). We performed Bayesian modeling to determine whether the relationship was significant, with reaction time as the dependent variable and feedforward population spiking response as the independent variable (Figure 1M). This analysis revealed significant predictive value in the independent variable’s (i.e., feedforward oddball response) coefficient estimates (*r*: M= -0.73, 89% CI=[-0.71, -0.74]; *β*: M=4.90, 89% CI=[4.68, 5.15]), explaining a large fraction of the variance (R^2^=0.62).

### Spatial evidence for feedforward selection

We next evaluated spatial evidence. The canonical columnar microcircuit puts forward layer-specific activations for feedforward vs. feedback computations (Bastos et al., 2012; Douglas and Martin, 1991; Douglas et al., 1989). These patterns are robustly observed in sensory cortex (Ferro et al., 2021; Mitzdorf, 1985; Nandy et al., 2017; Pettine et al., 2019; Schroeder et al., 1998; van Kerkoerle et al., 2017, 2014; Westerberg et al., 2022, 2021) (Figure 2A). Differences in granular layer (L4) synaptic activation as a function of oddball vs. distractor stimulation thus would indicate feedforward oddball signaling. Analyzing the fastest reaction time trials, we indeed observed a significant difference in synaptic activity L4 as a function of oddball vs. distractor presentations to the column’s population receptive field (Figure 2B). This result indicates differences at the level of feedforward input into V4. We quantified this relationship using the modeling techniques used for spiking data, listed above (Figure 2C). We again found a significant relationship between L4 feedforward synaptic activation to the attention-grabbing oddball and reaction time (*r*: M=-0.56, 89% CI=[-0.54, -0.57]; *β*: M=1.70, 89% CI=[1.64, 1.77]; R^2^=0.26).

**Figure 2.**
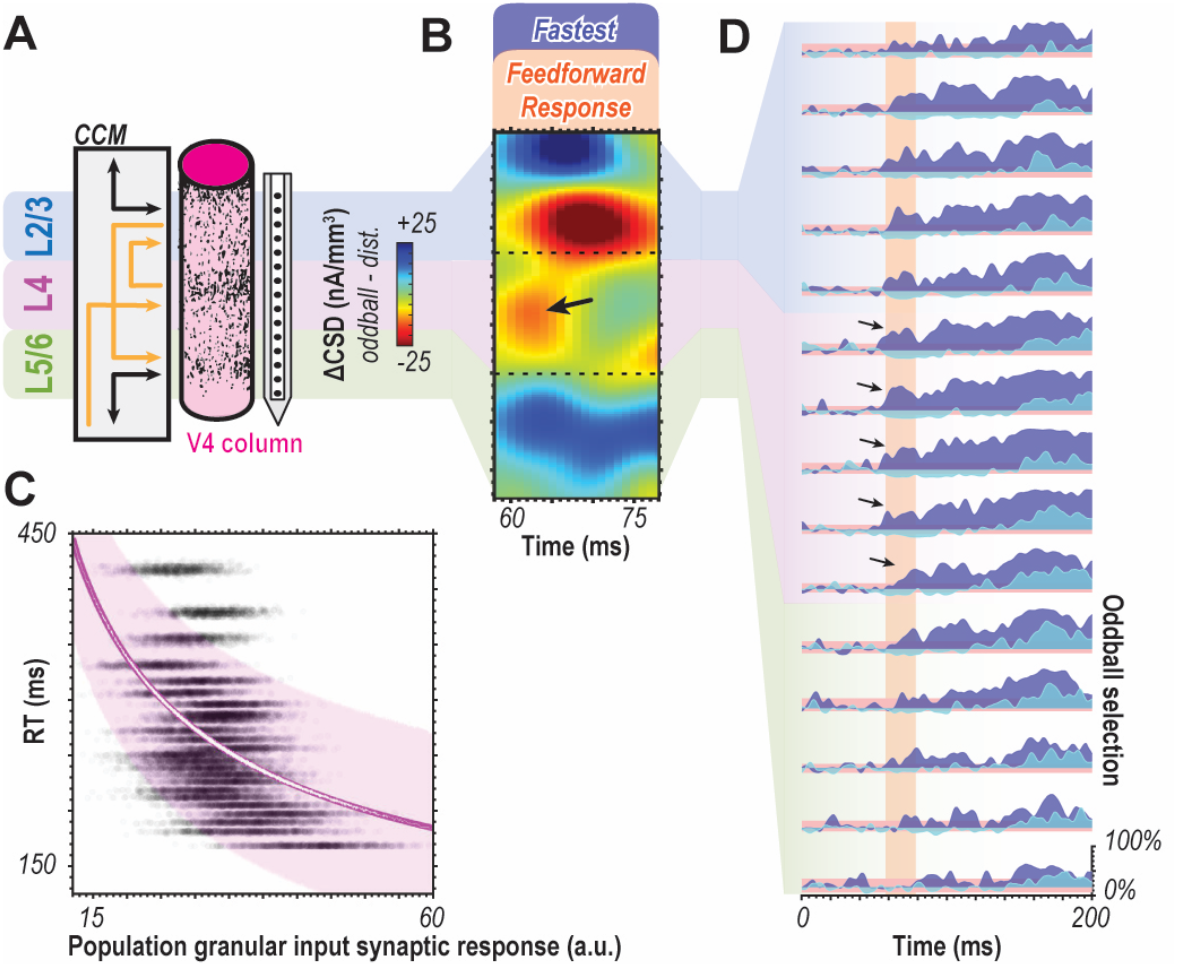
Spatial evidence for feedforward attentional selection across the canonical cortical microcircuit. **(A)** Cartoon illustrating the key laminar compartments (upper-L2/3, middle-L4, and deep-L5/6) with canonical microcircuit connectivity. Since earlier stages of sensory processing primarily project to L4 while feedback terminals innervate the layers above and below, feedforward computations should be indicated by differences in initial middle layer activation. **(B)** Laminar current source density (CSD, indicating synaptic activation) difference between oddball and distractor for the fastest, attention-capturing trials. Window shows 50 ms at the time of feedforward activation of the cortical column. Arrow indicates difference present in L4 where the oddball evoked a greater response. **(C)** Bayesian modeling of 24-bin data with population size 250 using synaptic currents in L4. Reaction time (RT) as a function of granular CSD magnitude (mean 58-78 ms post-display) to oddball, 1000 trials for each bin. Magenta line is the result of a power function fit. White data within magenta line are median estimates for simulated trials. Magenta cloud shows 89% credible interval of median estimates for fit. Slope indicates a negative relationship between L4 CSD magnitude to oddball stimulus and behavioral RT. **(D)** Oddball detection by cortical depth for fastest and slowest bins relative to visual display for population size 250 using multiunit responses. Samples were localized to each depth (n=15). Arrows denote significant oddball detection in the feedforward response in the middle layer. Orange highlights initial 20 ms window of feedforward visual response.

In further evaluating the laminar profile of oddball detection, we observed greater than chance detection during the initial response in the granular input layers for the fastest response trials (Figure 2D). Both findings indicate feedforward computation of attentional capture in sensory cortex. In other words, the oddball stimulus gets emphasized over other stimuli during the initial volley of synaptic activation following stimulus onset. The underlying computation thus happens either at this moment and location, prior to that, or both. If the oddball detection occurred at a previous (upstream) location of visual processing, it thus must have been derived without feedback from area V4 or other downstream areas. Also note that the initial activation of V4 input layers precedes full sensory activation of earlier areas, such as V1 and V2 (Schmolesky et al., 1998), as well as the onset of distinguishable feedback responses in these areas (van Kerkoerle et al., 2014). This context further suggests that pop-out oddball detection occurs within the first wave of stimulus-evoked spikes (VanRullen and Thorpe, 2002) rather than during reverberant processing. Summarizing these findings, we find both temporal and spatial evidence for feedforward generation of a priority signal for attentional capture.

### Errant selection precedes errant behavior

An interesting secondary question is whether attentional capture in the feedforward response is entirely a factor of salience or if the information is truly relayed as a priority signal (Fecteau and Munoz, 2006). While priority signals have been described for frontal (Thompson et al., 2005), parietal (Ipata et al., 2009), and temporal (Stemmann and Freiwald, 2019) cortex, there has been no strong neurophysiological evidence for sensory cortical priority signals. We thus decided to test for the presence of priority signals in sensory cortex. While incorrect behavioral responses are a small minority in pop-out search, the monkeys performed sufficient trials to yield a representative sample of error trials. We thus can determine whether the population signal in V4 reflects the salience of the stimulus (which is constant between correct and error trials) or priority (which differs between correct and error trials). Crucially, examined this process during the feedforward response period.

Two alternative hypotheses emerge from this line of reasoning. If the feedforward response reflects salience, we expect robust attentional capture for the oddball, even when a distractor was (erroneously) selected as the target. If the feedforward response computes priority, however, we expect the population response to reflect the incorrect target selection (Heitz et al., 2010; Thompson et al., 2005). We performed population reliability analysis to distinguish between these two possibilities.

In line with the hypothesis that the feedforward sensory response represents a priority signal, V4 population responses selected (misidentified) the distractor, errantly capturing attention (Figure 3A). Moreover, this selection was present in the initial response (Figure 3B).

**Figure 3.**
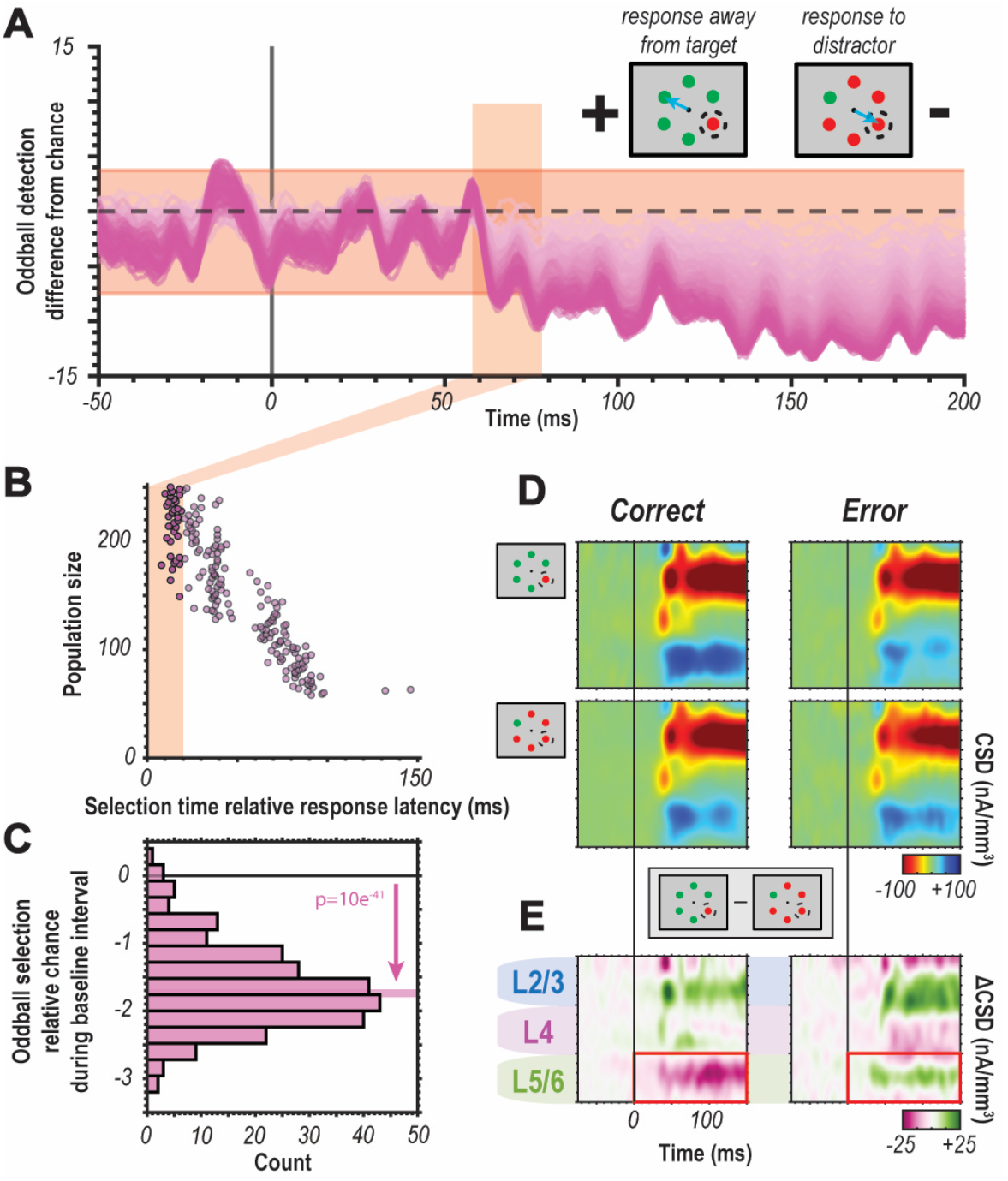
Errant selection precedes errant behavior. **(A)** Oddball selection for population size 1-250 (light to dark magenta) on incorrectly performed trials. Negative selection accuracy values indicate selection for the distractor. Feedforward visual response time window indicated in orange. Cartoon visualization of error types in upper righthand corner. **(B)** Selection times for prioritized distractor for each population size. Feedforward response time window highlighted in orange. **(C)** Histogram of oddball selection during baseline period (50 ms prestimulus window). Small, but reliable bias in selection for misprioritized distractor observable before visual display (result of two sample t test indicated in plot). **(D)** Comparison of synaptic activity for correct (left) vs. incorrect (right) trials across sessions for stimulus captured (top) vs. distractor (bottom) conditions. **(E)** Difference in synaptic activity between captured stimulus and distractor conditions for correct (left) vs. incorrect (right) trials. Red arrows indicate notable difference in granular input sink and red box denotes observed difference in deep-layer synaptic source.

Somewhat unexpectedly, we also found a small but significant selection bias during the pre-display (fixation) period (Figure 3C). This observation suggests errant capture could be partially explained by modulated ongoing activity. In further pursuing this question, we observed that errant capture was predominantly related to changes in synaptic activity in the deep layers of cortex (Figure 3D-E). Deep layers in V4 have been linked to behavioral output, and the difference found here is in line with that association (Pettine et al., 2019).

### Prestimulus modulation of pertinent feature selective columns

After noting the relationship between baseline activity and behavior in error trials (Figure 3), we hypothesized that coordinated modulation of baseline activity (Figure 4A-B) could bias capture. Previous work has implicated altered baseline activity in perceptual sensitivity to visual objects (Luck et al., 1997; Supèr et al., 2003; van der Togt et al., 2006; van Vugt et al., 2018). We thus structured the task to induce feature priming. Specifically, we employed “priming of pop-out” to promote attentional capture of a specific feature, such as the red or green color (Figure 4C). Searching for the same feature (e.g., red oddball among green distractors) repeatedly results in faster reaction times (Maljkovic and Nakayama, 1994). Swapping the target feature reinitiates priming for the new feature. This effect translates across species (Westerberg and Schall, 2021) and is also observed here (Figure 4D). Neural correlates of attentional priming exist in frontal (Bichot and Schall, 2002) and visual cortex (Westerberg et al., 2020); however, these findings do not explain how feature representations are promoted for capture. This latter type of attentional priming can be thought to reflect changes in the attentional prioritization of salient features.

**Figure 4.**
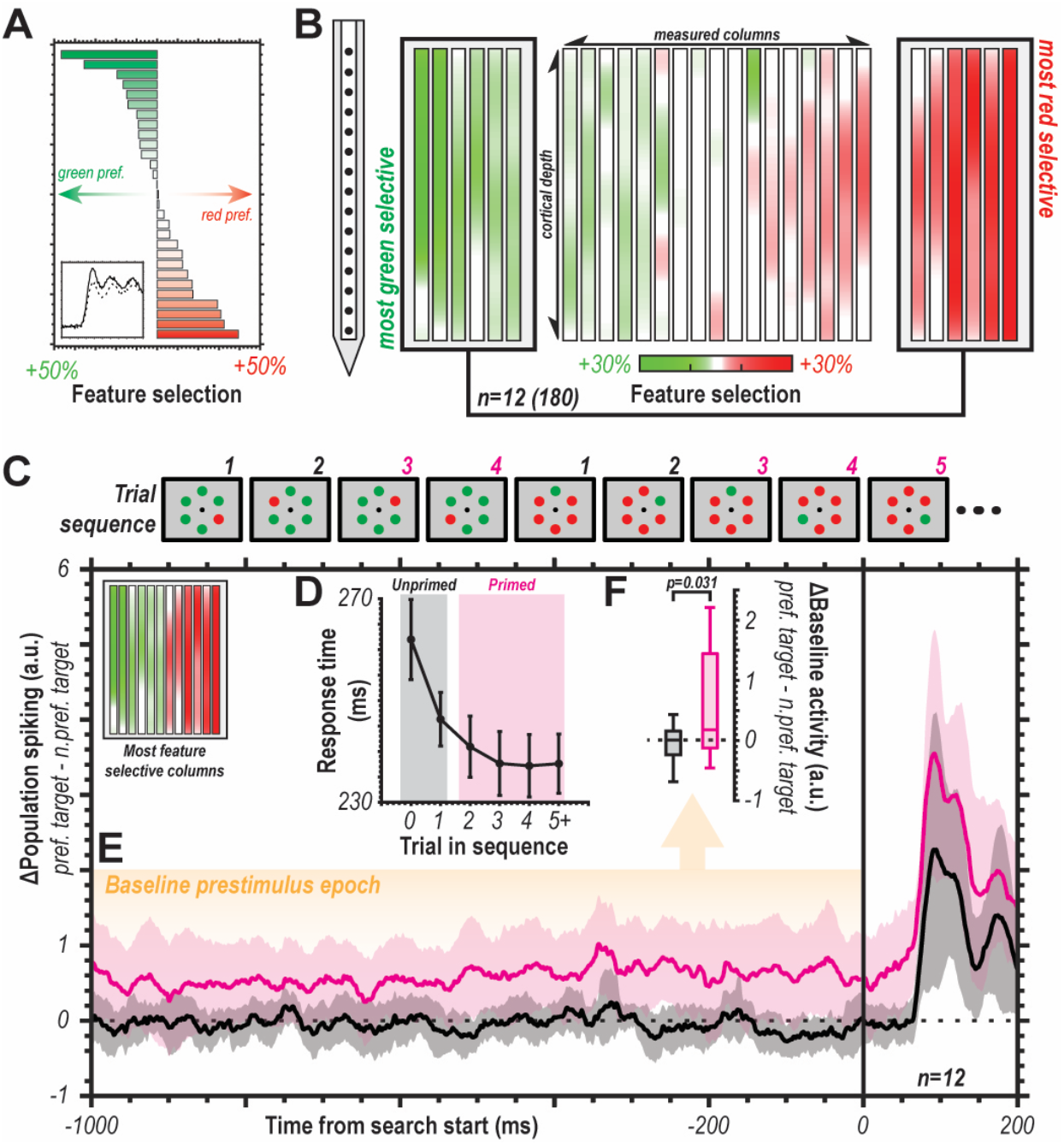
Selection history modulates ongoing activity. **(A)** Columnar feature selectivity organized from most green-preferring to most red-preferring columns (n=29). Bar size indicates degree of feature selectivity across each column’s multiunit spiking activity. Inset displays representative average red vs. green response for a red-preferring column (solid line, red response; dashed line, green response). Selectivity metric represents application of population reliability analysis at the individual multiunit level for selection of red vs. green. **(B)** Feature selectivity along depth for each column. Six most red- and green-preferring columns highlighted for further analysis. **(C)** Example trial sequence where target feature (red among green or vice versa) repeated before switching. This induced priming of target feature where primed (magenta) can be compared to unprimed responses (gray). **(D)** Reaction time (medians with 95% confidence intervals) decreases with repeated search for a feature (n=29 sessions). **(E)** Mean difference in baseline population spiking between preferred feature oddball and non-preferred feature oddball for unprimed (black) and primed (magenta) trials for feature columns (inset top-left) (n=12 columns, n=180 multiunits). Clouds are 95% confidence intervals. **(F)** Boxplot of differences in baseline spiking for preferred feature target vs. non-preferred feature target for unprimed vs. primed trials. Central mark indicates median, and the bottom and top edges indicate the 25^th^ and 75^th^ percentiles. Whiskers are +/-2.7 SD. Statistic is two-sided t test.

We tested this by first identifying color-selective feature columns in V4. Topographic organization for color exists across V4 (Conway and Tsao, 2009; Tanigawa et al., 2010; Westerberg et al., 2021) and can be observed at the cortical columnar level (Figure 4A-B). We measured the difference in baseline activity for the preferred vs. non-preferred feature before and after establishing behavioral relevance via priming (Figure 4E). We found that baseline activity was significantly higher in columns selective for a color when the color was behaviorally relevant (Figure 4F). Note that this difference in firing persists in the absence of visual stimulation, thereby supporting the proposition of feedback modification in the case of task repetitions (priming).

### Feedforward selection does not require priming

With the understanding that feature-selective cortical columns can be modulated to promote their responses for attentional selection, we sought to determine whether feedforward selection was only present under priming conditions. To test this, we reduced the data to trials immediately following a switch in the priming sequences (e.g., the trial when the task switched from red among green to green among red). In these trials, the search target is not primed (and may in fact be negatively primed). We then repeated the Bayesian modeling of reaction time as predicted by oddball multiunit responses for these unprimed trials. The procedure was identical to that described in Figure 1. We found significant predictive value in the independent variable’s coefficient estimates (*r*: M=-0.45, 89% CI=[-0.43, -0.48]; *β*: M=1.73, 89% CI=[1.64, 1.86]) explaining a sizeable fraction of the variance (R^2^=0.18). This finding indicates that even when feedback is putatively promoting the feature representations associated with the distractors, the predictive relationship between oddball feedforward responses and reaction times is preserved. Thus, even though presentation of an oddball array leads to modulating feedback following task completion, this feedback cannot sufficiently explain the feedforward selection we observed. In other words, feedforward attentional selection during pop-out is not predicated on prior feedback.

## Discussion

We found that the magnitude of neuronal population responses in sensory cortex to an attention-grabbing stimulus is predictive of reaction time and accuracy during attentional capture. Crucially, this relationship emerged in the earliest parts of the stimulus-driven response propagating feedforward. Moreover, oddball detection was observed in the initial current sink that marks the synaptic activity propagating through the granular input layer of sensory cortex. Remarkably, feature columns were tonically modulated by repeated task demands, adjusting ongoing activity to promote attentional capture for consistently pertinent features. In line with this notion, errantly biased activation during the pre-stimulus epoch was associated with errant behavior. These findings demonstrate an unexpectedly dominant role for sensory cortex in dictating attentional priority. From a theoretical perspective, these observations resolve a long-standing debate on attentional capture (Folk et al., 1992; Luck et al., 2021; Theeuwes, 1993). Specifically, we find that a priority signal is automatically generated in a feedforward fashion. That signal is then used for tonic modulation via feedback to promote the detection of similar objects in subsequent searches. In other words, our attention is automatically captured by salient features. The speed of our behavior in response to these objects is dictated by the variability in their engendered sensory response. However, behavioral goals and historical context can influence which features we are more sensitive to, effectively promoting repeated attentional capture for objects comprised of those features.

In evaluating population codes for the representation of sensory information, it becomes important to consider the size of a neural population that allows for certain computations to be performed (Averbeck et al., 2006). For example, in behavioral tasks, an observation requiring an inordinately large population of neurons, could be inconsequential if that information cannot be relayed downstream for the execution of behavior. In our study, gaze must be redirected for our behavioral measure (reaction time) to occur. This process likely engages areas like the frontal eye fields (FEF) (Schall, 2015, 2004), which receive sensory information from V4 (Anderson et al., 2011; Gregoriou et al., 2012; Ninomiya et al., 2012; Schall et al., 1995; Stanton et al., 1995; Ungerleider et al., 2008; Zhou and Desimone, 2011). Therefore, the population representation of the oddball stimulus must be relayed effectively from populations of V4 to FEF neurons through biologically plausible pathways. With this in mind, it is worth noting that reliable oddball detection at the population level can be observed in sets of 20 or fewer multiunits during the feedforward period (Figure 1, Extended Data Figure 3).

It is also interesting to consider population size as a source for the variability of reaction time that is not explained by the modeling. The different percentages of time the oddball is selected between traces suggest that the exact population size does impact the magnitude of detectability, at least in this population reliability metric. Therefore, it could be hypothesized that some variability observed in the behavioral response as a function of behavioral capture might be due to the size of the population propagating the signal.

While we have found that modification of the priority signal exists tonically, is there an antecedent? That is, what instigates the persistent change in activity found in the behaviorally relevant feature columns? One hypothesis is that the initial presentation of the attention-grabbing oddball leaves sensory cortex in an altered state, more sensitive to the established pertinent feature (Westerberg and Schall, 2021). Adaptation in sensory cortex can have potent effects (Albrecht et al., 1984; Carandini, 2000; Clifford et al., 2007; Dragoi et al., 2000; Kohn, 2007; Kohn and Movshon, 2004), and is implicated in changing response characteristics at the level of cortical columns (Hansen and Dragoi, 2011; Westerberg et al., 2019). An alternative view might be that the frontal cortex regulates feature-based attentional modulation in the visual cortex; a candidate area (VPA) has been identified that could serve as a source (Bichot et al., 2019, 2015). Either hypothesis does not change the interpretation that the sensory cortex automatically computes attentional capture; however, resolving between them would provide insight into the minimum required neural circuitry to modify the priority signal.

## Methods

### Experimental Model and Subject Details

#### Animal Care

Two male macaque monkeys (*Macaca radiata*; monkey Ca, He) participated in this study. All procedures were in accordance with the National Institutes of Health Guidelines and the Association for Assessment and Accreditation of Laboratory Animal Care International’s Guide for the Care and Use of Laboratory Animals, and approved by the Vanderbilt Institutional Animal Care and Use Committee in accordance with United States Department of Agriculture and United States Public Health Service policies. Animals were pair-housed. Animals were on a 12-hour light-dark cycle and all experimental procedures were conducted in the daytime. Each monkey received nutrient-rich, primate-specific food pellets twice a day. Fresh produce and other forms of environmental enrichment were given at least five times a week.

#### Surgery

All surgical procedures were performed under aseptic conditions. Anesthesia was conducted with animals under N_2_O/O_2_, isoflurane (1-5%) anesthesia mixture. Vital signs were monitored continuously. Expired PCO_2_ was maintained at 4%. Postoperative antibiotics and analgesics were administered while animals remained under close observation by veterinarians and staff. Monkeys were implanted with a custom-design head post and MR-compatible recording chamber using ceramic screws and biocompatible acrylic. A craniotomy over V4 was opened concurrent with the recording chamber.

### Method Details

#### Magnetic resonance imaging

MR images for chamber localization and guiding of linear electrode penetrations perpendicular to the cortical surface were taken from anesthetized animals placed inside a 3T MRI scanner (Philips). T1-weighted 3-dimensional MPRAGE scans were acquired with a 32-channel head coil equipped for SENSE imaging. Images were acquired using a 0.5 mm isotropic voxel resolution with the following parameters: repetition time 5 s, echo time 2.5 ms, and flip angle 7°.

#### Identification of V4

Recordings took place on the convexity of the prelunate gyrus in approximately the dorsolateral, rostral aspect of the V4 complex, where receptive fields are located at about 2–10 degrees of visual angle (dva) eccentricity in the lower contralateral visual hemifield (Gattass et al., 1988). Laminar recordings took place at locations where the array could be positioned orthogonal to the cortical surface, as verified by MRI and neurophysiological criteria (i.e., overlapping receptive fields). Recording sites were also confirmed via histological staining by dipping the electrode arrays in diiodine prior to the final recordings in monkey He (Westerberg et al., 2020).

#### Task design: Pop-out search

Monkeys viewed arrays of stimuli presented on a CRT monitor with 60 Hz refresh rate, at 57 cm distance. Stimulus presentations and task timing was controlled using TEMPO (Reflective Computing). Visual presentations were monitored with a photodiode positioned on the CRT monitor so that electrophysiological signals could be reliably aligned offline. Red (CIE coordinates: x=0.648, y=0.331) and green circles (CIE coordinates: x=0.321, y=0.598) were used as stimuli, rendered isoluminant to a human observer at 2.8 cd/m^2^ on a uniform gray background. As we are limited to two colors and cannot account for potential differences in perceived brightness between macaques, we qualify our two stimuli as distinct ‘features’ at the intersection of color and luminance information. Nonetheless, we report the colors used in this study for the ideal human observer. Cone excitation was computed from the CIE coordinates and luminance (Cole and Hine, 1992). The following cone excitations were measured for red: *ε*_*L*_ =2.37, *ε*_*M*_ =0.43, *ε*_*S*_ =0.0014; green: *ε*_*L*_=1.74, *ε*_*M*_ =1.06, *ε*_*S*_=0.0030; and the background: *ε*_*L*_ =1.86, *ε*_*M*_ =0.94, *ε*_*S*_ =0.023. Cone contrasts for red stimulus were found to be: *C*_*L*_ =0.27, *C*_*M*_ =-0.54, *C*_*S*_ =-0.94; and the green stimulus: *C*_*L*_ =-0.06, *C*_*M*_ =0.13, *C*_*S*_ =-0.87.

Trials were initiated when monkeys fixated within 0.5 dva of a small, white fixation dot (diameter = 0.3 dva). The time between fixation acquisition and array presentation varied between 750–1250 ms, taken from a nonaging foreperiod function to eliminate any potential effect of stimulus expectation (Näätänen, 1971, 1970; Nickerson and Burnham, 1969). Following the fixation period, the stimulus array consisting of 6 items was presented. Stimuli were scaled with eccentricity at 0.3 dva per 1 dva eccentricity so that they were smaller than the estimated V4 receptive field size (0.84 dva per 1 dva eccentricity (Freeman and Simoncelli, 2011)). The polar angle positioning of the items relative to fixation varied from session to session so that one item of the stimulus array was positioned at the center of the population receptive field under study. Items were spaced such that only one item was in the V4 receptive field, with uniform spacing in polar angle and equal eccentricity.

Monkeys engaged in a search task while viewing the stimulus array. One item in the array was a different feature (red or green, respectively) from the others. Position of the oddball on each trial was randomly chosen with equal probability for any of the positions (16.6%). Monkeys earned fluid reward for shifting gaze directly to the oddball item within 1000 ms of array presentation and maintaining fixation within a 2–5 dva window around the oddball for 500 ms.

Eye movements were monitored continuously at 1 kHz using an infrared corneal reflection system (SR Research). If the monkey failed to look at the oddball, no reward was given, and a 1-5 s timeout ensued. Trials were organized into blocks such that the animal searched for the same target feature for 5-15 repetitions. Target feature remained the same, but target location varied randomly. Completing the block resulted in the target and distractor features swapping.

#### Neurophysiology

Laminar extracellular voltages were acquired at 24.4 kHz resolution using a 128-channel PZ5 Neurodigitizer and RZ2 Bioamp processor (Tucker-Davis). Raw signals were output between 0.1 Hz and 12 kHz. Data were collected from 2 monkeys (left hemisphere, monkey Ca; right hemisphere, He) across 70 recording sessions (n=31, monkey Ca; n=39, He) using 32 channel linear microelectrode arrays with 0.1 mm interelectrode spacing (Plexon). Each recording session, electrode arrays were introduced into the prelunate gyrus through the intact *dura mater* using a custom micromanipulator (Narishige). Electrode arrays were positioned so they spanned all layers of V4 and had a subset of electrodes positioned outside of cortex. 29 (n=20, monkey Ca; n=9, He) of 70 sessions were included in the final analysis. The remaining 41 sessions were found to either not have a discernable CSD profile for laminar alignment, not be orthogonal to the cortical surface, or not have enough priming blocks for the feedback mechanism analysis and were thus removed from analysis.

#### Receptive field mapping

To determine the orientation and eccentricity of the visual receptive fields, monkeys performed a receptive field mapping task prior to the main task. Monkeys fixated for 400–7000 ms while a series of 1–7 stimuli were presented that spanned the visual field contralateral to the recording chamber. Stimuli were 5 high-contrast concentric white and black circles that scaled in size with eccentricity (0.3 dva per 1 dva eccentricity). In all recording sessions, stimuli could appear in a random location. These random locations spanned the lower visual quadrant contralateral to the recording chamber. Location spacing was in 5° angular increments relative to fixation and in eccentricities ranging from 2 dva to 10 dva in 1 dva increments. Each stimulus was presented for 200–500 ms with an interstimulus interval of 200–500 ms. If the animal maintained fixation for the duration of the stimulus presentation sequence, they received a juice reward. During this receptive field mapping task, multiunit activity, gamma power (30-90 Hz), and evoked local field potentials (LFPs, 1–100 Hz) were measured across all recording sites. Online, we measured the response across visual space for each recording site. Recordings proceeded to the feature search task if there was qualitative homogeneity of receptive fields along depth. Receptive field overlap for these data have been reported previously (Westerberg et al., 2021). The receptive field center was chosen to be the stimulus location that evoked the largest response along the depth of recording sites. Following receptive field identification, the stimulus array in the feature search task was then oriented so that its eccentricity coincided with the location of the receptive field (eccentricity: 3–10 dva) and a single array item was placed at the center of the receptive field (size: 0.9–3 dva).

#### Identification of cortical laminae

Positions of the individual recording sites relative to the layers of V4 were determined using current source density (CSD) analysis. CSD reflects local synaptic currents (net depolarization) resulting from excitatory and inhibitory postsynaptic potentials (Mitzdorf, 1985). CSD was computed from the raw neurophysiological signal by taking the second spatial derivative along the electrode contacts (Maier et al., 2011, 2010; Nicholson and Freeman, 1975; Schroeder et al., 1998; Westerberg et al., 2019). CSD activation following presentation of a visual stimulus reliably produces a specific pattern of activation which can be observed in primate visual cortex (Maier et al., 2010; Schroeder et al., 1998), including V4 (Bollimunta et al., 2008; Givre et al., 1994; Nandy et al., 2017; Westerberg et al., 2022, 2021). Specifically, current sinks following visual stimulation first appear in the granular input layers of cortex, and then ascend and descend to the extragranular compartments. To compute the CSD from the LFP, we used previously described procedure (Nicholson and Freeman, 1975):

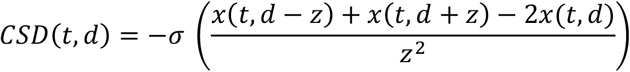

where the CSD at timepoint t and at cortical depth *d* is the sum of voltages *x* at electrodes immediately above and below (*z* is the interelectrode distance) minus 2 times the voltage at *d* divided by the interelectrode-distance-squared. That computation yields the voltage local to *d*. To transform the voltage to current, we multiplied that by *-σ*, where *σ* is a previously reported estimate of the conductivity of cortex (Logothetis et al., 2007). For each recording session, we computed the CSD and identified the initial granular layer (L4) input sink following visual stimulation. Sessions were aligned using the bottom of the initial feedforward input sink as a functional marker. We defined the size of individual laminar compartments uniformly relative to space. Throughout, ‘middle’ refers to the estimate of the granular input layer 4 (0.5 mm space above the CSD initial sink functional marker), ‘upper’ refers to the estimate of supragranular layers 2 and 3 (0.5 mm space above the L4 compartment), and ‘lower’ refers to the estimate of infragranular layers 5 and 6 (0.5 mm space below the L4 compartment).

#### Population spiking

Spiking activity at the level of multiunits was used for control analyses as it reliably reflects neural population dynamics (Trautmann et al., 2019). Detection of multiunit activity was achieved through previously described means (Legatt et al., 1980). This method has proved useful across brain areas and research groups (Logothetis et al., 2001; Self et al., 2013; Shapcott et al., 2016; Supèr and Roelfsema, 2005; Teeuwen et al., 2021; Tovar et al., 2020; Westerberg et al., 2020). Briefly, broadband neural activation was filtered between 0.5-5 kHz, the predominate range of spiking activity. The signal was then full-wave rectified and filtered again at half the original high-pass filter (0.25 kHz) thereby estimating the power of the multiunit activity. For filtering, we used a 4^th^-order Butterworth filter. Spiking responses were baseline corrected by subtracting the average activity in the 100 ms window preceding visual display onset at the trial level. This baseline correction was not performed for the feedback analysis.

#### Feedforward sensory response window

Determining the implications of the feedforward response to attentional capture required accurate identification of the timing of said feedforward response. Here, the window of the feedforward response is defined as the 20 ms following the time at which the mean population spiking response first reaches 50% of its maximum response. This definition yielded a response latency of 58 ms (mean=58, 95% confidence interval=[57, 59]) with the window being defined as 58-78 ms following visual display onset.

### Quantification and Statistical Analysis

#### Sorting responses by reaction time

Several analyses were conditioned on sorting trials by behavioral reaction time. Through this procedure, trials were rank ordered by reaction time from fastest to slowest on a session-by-session basis. For example, if session *n* contained 2000 trials, each trial was ranked from 1 to 2000 by reaction time, and then normalized as a percentile. This ranking was completed individually for each session so that individual sessions could be sampled equally for each binned condition. Two binning procedures were performed, one coarse (4 bins) and one fine (25 bins). For the fine-binning-conditioned analyses, the slowest bin (slowest 2% of trials) was omitted from analysis as outliers in otherwise efficient, pop-out search.

#### Population reliability analysis

Population reliability analysis was used to establish whether and when populations of V4 neurons selected an attention-grabbing oddball (Bichot et al., 2001). Crucially, this analysis estimates when selection occurs in time as well as how many neural units are required for this selection to occur reliably. This analysis is performed by simulating trials using data from the entire population of multiunit responses across all sessions. Each simulated trial is defined as an event where a behavioral response must be made to a stimulus with multiple alternatives present. For this computation, each alternative is represented by the response of a distinct neural population with a predetermined size. We varied the population size between 1 and 250. In this study, the search task contains six alternatives, thus we estimated a population response for each of the six stimuli. The population response is defined as the sum of responses of each of the sampled responses – where each response is an empirically measured trial-level multiunit response to the stimulus germane to the alternative’s response that is being estimated. Therefore, we chose five distinct, randomly sampled population responses to distractor ‘alternatives’ and one to an oddball ‘alternative’. For each point in time within each simulated trial, we measured which alternative provoked the highest response magnitude. This selection metric represents our estimated priority signal. By simulating more and more trials (n=1000 for each computation in this study), we computed the percent of time each alternative is selected at each timepoint during a simulated trial. Here, we were specifically interested in the percent of time the oddball was selected by this measure. We defined this percentage as the selection accuracy in identifying the behaviorally relevant oddball. For six objects, chance (selection invariability between population responses) was calculated to be 16.67%. We computed empirical selection bounds to estimate when the selection accuracy exceeded chance by measuring the variability in selection accuracy for all 1–250-unit populations during the baseline, prestimulus display, epoch and setting the thresholds to the 99% confidence interval.

#### Bayesian modeling

Bayesian modeling was performed using Stan through the RStan interface. Sampling was done using Markov chain Monte Carlo (MCMC) methods with the following parameters: chains, 4; warmup samples, 2000; total samples, 5000; thinning, 2. A power function fit was assessed. The outcome *RT*_*i*_ (reaction time on simulated trial *i*) can be modeled as:

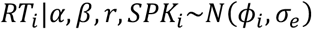

where:

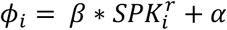

Reaction time (*RT*) for simulated trial *i* is modeled as the population multiunit spiking activity (*SPK*) for simulated trial *i* with coefficient *β. r* is the exponent for the power function fit. Population spiking activity was defined as the sum of activity across the population. In supplementary analyses we also explored the same relationship to reaction time, but with the magnitude of the granular input sink taken as the average of 5 recording channels immediately above the L4/5 boundary for each trial. We set minimally informative priors for the power function fit as: *α*∼*LogNormal*(0,0.5), *β*∼*LogNormal*(1,0.5), *r*∼*Gamma*(1,3), *σ*_*e*_∼*Gamma*(0.5,5).

From this modeling, we computed median estimates for each simulated trial as well as the associated 89% credible intervals. We also computed reaction time estimates for the range of population spiking responses observed using the median estimate of each coefficient. In evaluating the posterior distributions, we were interested in the median (M) estimates for the coefficients *β* and *r* as they reflect the predictive utility of the independent variable (population spiking response). In particular, non-zero *β* and *r* different than 1 indicate significant utility provided the 89% credible intervals (89% CI) for those estimates do not include their respective non-predictive values.

#### Feature selectivity

Feature (red vs. green) selectivity was derived from population spiking observed along recording sites. Responses were taken when a red stimulus was presented to the receptive field of the cortical column and when a green stimulus was present in the receptive field. We employed a two-alternative version of the population reliability analysis to estimate the selectivity of each multiunit for red vs. green. For each multiunit we took 100 red and 100 green stimulus presentations (effectively population size 100) 1000 times (bootstrapped simulated trials) for the reliability analysis. Specifically, we took the average response 60-160 ms following visual display. This yielded a selection accuracy metric for red vs. green where deviation from 50% chance indicated preference for one color or the other. Presence of feature selective columns in this dataset was confirmed in previous reports (Westerberg et al., 2022, 2021).

## Acknowledgments

The authors would like to thank E. Sigworth for their technical support and J. Theeuwes, G. Logan, T. Palmeri, G. Cox, S. Lilburn, and G. Bahg for their comments. This work was supported by the National Eye Institute [grant numbers: R01EY027402, R01EY019882, R01EY008890, P30EY008126]. J.A.W. was supported by fellowships from the National Eye Institute [grant numbers: F31EY031293, T32EY007135]. Imaging support was provided by the Vanderbilt University Institute for Imaging Science through a grant from the National Institutes of Health Office of the Director [grant number: S10OD021771]. Supercomputing resources were provided by the Vanderbilt University Advanced Computing Center for Research and Education.

## Author contributions

Conceptualization, J.A.W, A.M., J.D.S., G.F.W.; Data Collection, J.A.W.; Formal Analysis, J.A.W.; Data Visualization, J.A.W.; Original Draft, J.A.W.; Revisions and Final Draft, J.A.W., A.M., J.D.S., G.F.W.

